# A conserved cell-type gradient across the human mediodorsal and paraventricular thalamus

**DOI:** 10.1101/2024.09.03.611112

**Authors:** Anton Schulmann, Ningping Feng, Pavan K Auluck, Arghya Mukherjee, Ruchi Komal, Yan Leng, Claire Gao, Sarah K Williams Avram, Snehashis Roy, Ted B Usdin, Qing Xu, Vesna Imamovic, Yash Patel, Nirmala Akula, Armin Raznahan, Vilas Menon, Panos Roussos, Laramie Duncan, Abdel Elkahloun, Jatinder Singh, Michael C Kelly, Michael M Halassa, Samer Hattar, Mario A Penzo, Stefano Marenco, Francis J McMahon

## Abstract

The mediodorsal thalamus (MD) and adjacent midline nuclei are important for cognition and mental illness, but their cellular composition is not well defined. Using single-nucleus and spatial transcriptomics, we identified a conserved excitatory neuron gradient, with distinct spatial mapping of individual clusters. One end of the gradient was expanded in human MD compared to mice, which may be related to the expansion of granular prefrontal cortex in hominids. Moreover, neurons preferentially mapping onto the parvocellular division MD were associated with genetic risk for schizophrenia and bipolar disorder. Midbrain-derived inhibitory interneurons were enriched in human MD and implicated in genetic risk for major depressive disorder.

## Main text

The mediodorsal thalamus (MD) is a large thalamic nucleus with strong reciprocal projections to the prefrontal cortex and is involved in many executive functions such as working memory, attention, and decision-making^1,2^. Dorsal midline structures adjacent to MD, the parataenial (PT) and paraventricular thalamic nuclei (PVT), have strong projections to medial prefrontal cortex, ventral striatum, and amygdala, and are involved in emotion regulation and motivation^3^. Converging evidence from volumetric, connectivity, and stroke lesion studies suggests that MD and its neighbors play important roles in psychiatric disorders, particularly psychosis^4–7^.

Despite their importance for cognition and psychopathology, the cell-type composition of MD and adjacent midline structures has not been completely characterized. Previous work in adult mice identified a cross-modal axis of thalamic neurons defined by distinct transcriptional, functional, and morphological properties^8^. This thalamus-wide gradient can be divided into three subdivisions (primary-secondary-tertiary), is conserved across species, and maps onto a thalamocortical connectivity gradient identifiable by diffusion tractography in humans^9^. Recent transcriptomic studies suggest that thalamus-wide cell-type gradients emerge during early prenatal development in mice and humans^10,11^. Transcriptomic and circuit dissection studies indicate that mouse MD and PVT harbor multiple transcriptionally and functionally distinct cell types^8,12–14^. However, their human correlates remain unknown.

To identify cell types in human MD and adjacent midline nuclei, we performed deep single-nucleus RNA-sequencing (snRNA-seq) on neurotypical controls from NIMH’s Human Brain Collection Core (HBCC, n=6) and Mount Sinai’s NIH Brain Tissue Repository (MSSM, n=4). MD was dissected from coronal slabs based on anatomical landmarks, followed by nuclei isolation and snRNA-seq via the 10X Chromium platform (Fig. 1a). Data underwent extensive quality control and filtering, clustering, and cell-type annotation based on marker genes and data from the Human Brain Cell Atlas^15^. Despite differences in tissue processing between the brain banks, both datasets showed the same major cell classes (Fig. 1b, Supplementary fig. 1a-d, see Methods for details). We identified excitatory neurons, marked by expression of *SLC17A6* (VGluT2), inhibitory neurons, which comprised nearly 40% of neurons in our samples and expressed *SOX14 –* consistent with midbrain origin – and several non-neuronal populations (Fig. 1c; Supplementary data table 1).

**Fig. 1.**
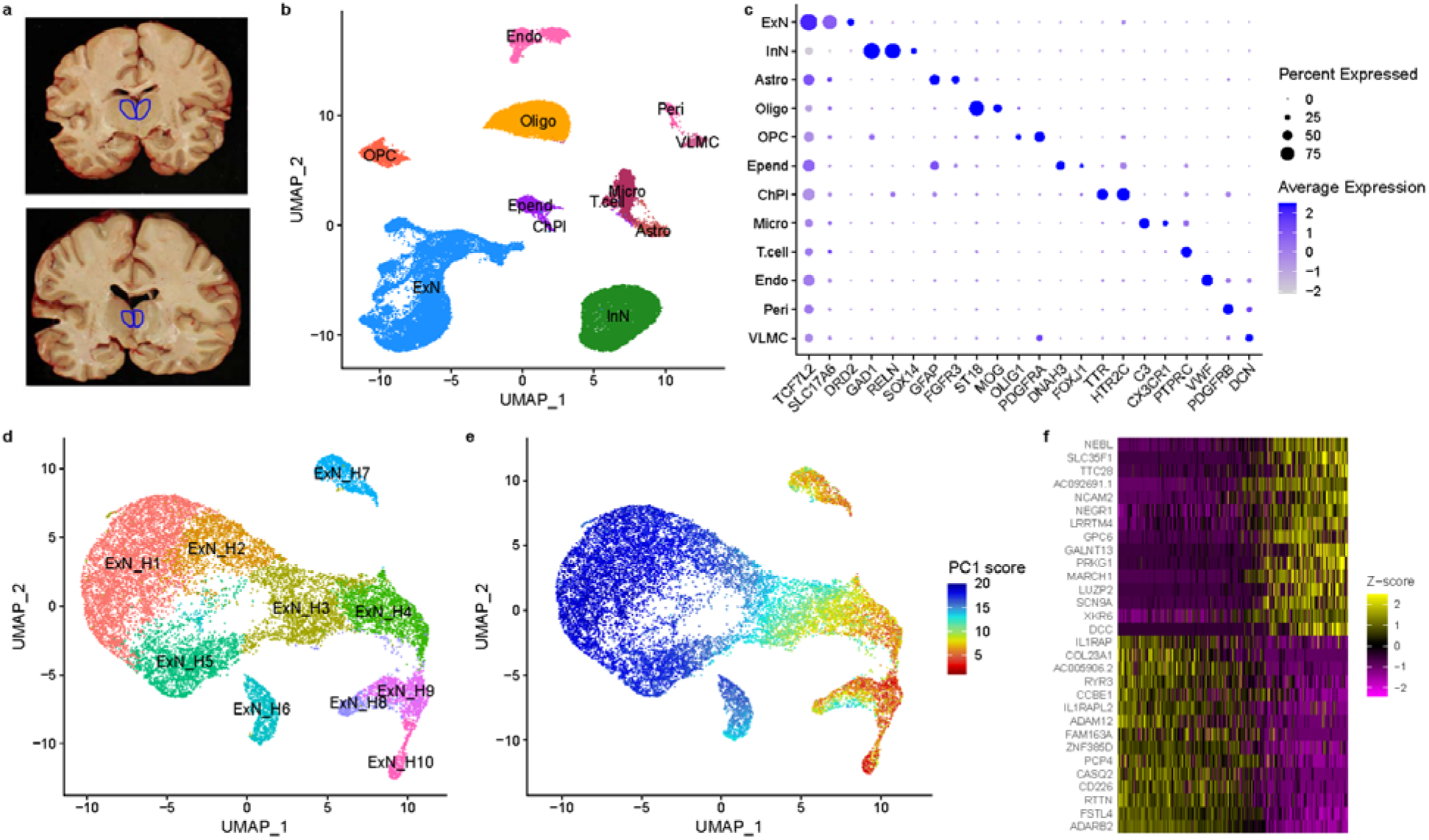
A cell type atlas of human MD and dorsal midline thalamus. a. Dissection approach targeting the MD and adjacent midline structures based on gross anatomical landmarks, illustrated by two example photographs of coronal slabs with region of interest highlighted in blue. b. UMAP plot illustrating annotation of major cell classes in human MD and midline thalamus (Excitatory Neurons [ExN]; Inhibitory Neurons [InN]; Astrocytes [Astro]; Oligodendrocytes [Oligo]; Oligodendrocyte Precursor Cells [OPC]; Ependyma [Epend]; Choroid Plexus [ChPl]; Microglia [Micro]; T-cells; Endothelial cells [Endo]; Pericytes [Peri]; Vascular leptomeningeal cells [VLMC]). See supplementary data table 1 for full list of marker genes. c. Dot plot illustrating expression pattern of marker genes associated with major cell classes shown in (b). (b) and (c) are based on HBCC samples; for MSSM see Supplementary fig. 1. d. UMAP plot of excitatory neurons, colored by cluster identities. e. UMAP plot of excitatory neurons colored by first principal component (PC1) score. f. Heatmap of top 15 genes with highest and lowest loadings on PC1. Cells are ordered by their PC1 score. (d), (e), and (f) are based on integration between HBCC and MSSM samples; for details, see Methods and Supplementary fig. 1.

To investigate thalamic projection neuron subpopulations in MD, we re-clustered the excitatory neurons, integrating HBCC and MSSM data (Fig. 1d, Supplementary fig. 1e-f). We found that most excitatory neurons in MD were part of a transcriptional gradient captured by the first principal component (PC1, Fig. 1e). Genes aligned with PC1 overlapped with and correlated in their loadings with those of a previously described trans-thalamic axis separating primary, secondary and tertiary thalamic cells in mice^8^ (Supplementary fig. 1g-h). Thus, projection neurons in human MD and midline thalamus exhibit a gene expression gradient that closely resembles a cell-type gradient with distinct functional and morphological properties in mice.

Human MD has traditionally been divided into a magnocellular (MDmc, medial), parvocellular (MDpc, lateral-ventral) and multiform/densocellular (MDdc, lateral-dorsal) part^16^. We hypothesized that these cytoarchitectonic subdivisions could correspond to transcriptionally defined neuronal populations. To map cell populations identified by snRNA-seq onto anatomical space, we applied spatial transcriptomics to two coronal thalamic sections, then deconvoluted the cell types, using our snRNA-seq data as a reference^17^. This revealed prominent spatial separation of neuronal populations. Some excitatory neuron clusters preferentially mapped to MDmc, MDpc, or adjacent nuclei, while inhibitory interneurons appeared to be particularly enriched in MDpc and MDdc (Fig. 2a; Supplementary fig. 2a).

**Fig. 2.**
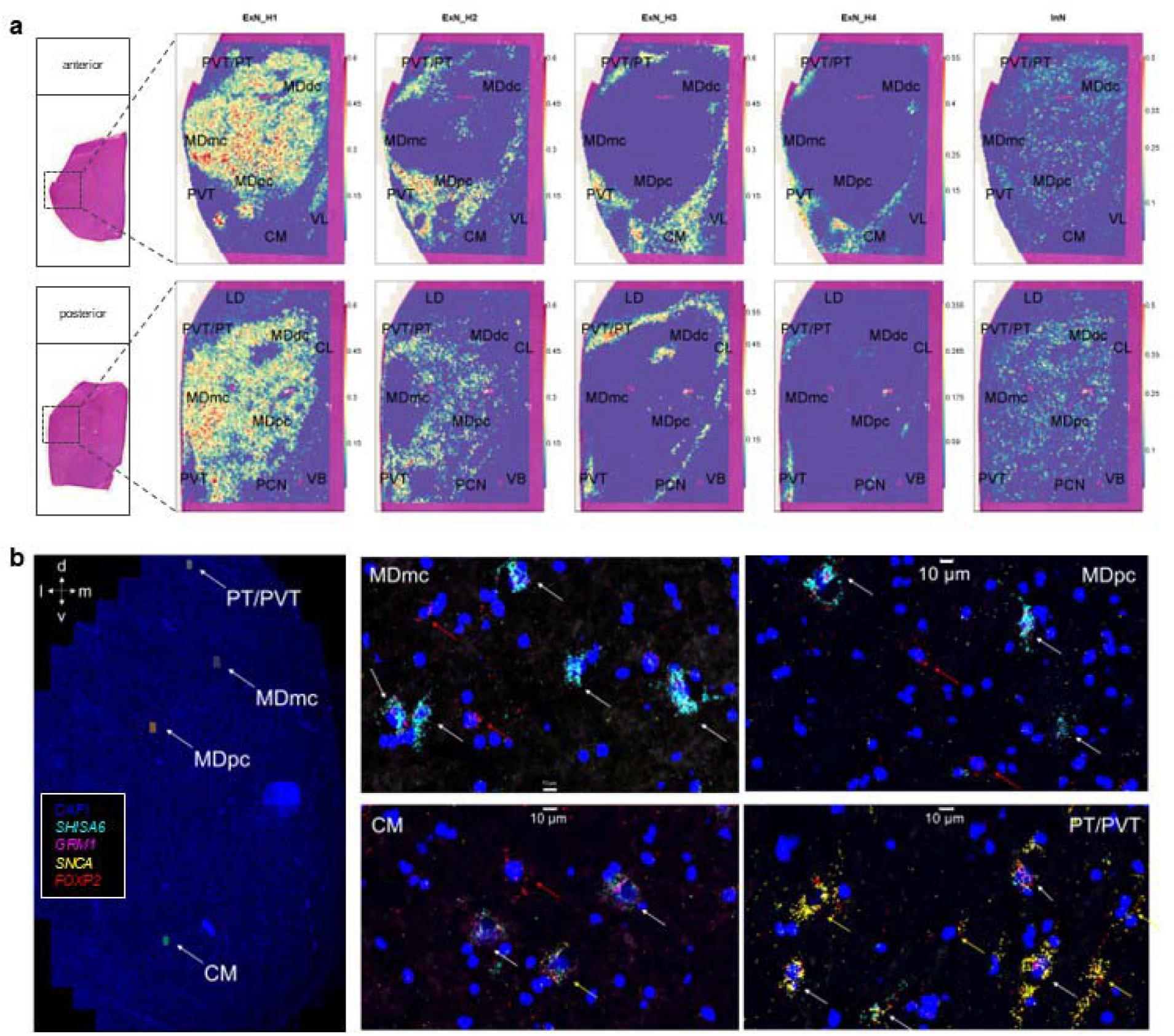
Spatial distribution of neuronal populations in human MD and midline thalamus. a. Spatial mapping of inferred cell type proportions for four excitatory neuron populations (first four columns) and thalamic inhibitory neurons (last column) across two coronal thalamic sections. Schematic (left) shows position of the 11×11mm wide capture area. Clusters shown preferentially mapped onto MDmc (ExN_H1), MDpc (ExN_H2), intralaminar nuclei, PT and PVT (ExN_H3, ExN_H4). Thalamic inhibitory neuron (InN) were most abundant in MDpc and MDdc. Major findings are summarized in Supplementary table 1. b. RNA-FISH of four marker genes (Primary markers: *SHISA6* and *GRM1*; Secondary/Tertiary markers *FOXP2* and *SNCA*) are shown across panels representative of four (sub)nuclei. MDmc and MDpc preferentially express primary markers, while CM and PVT/PT express tertiary markers with intermediate cells co-expressing markers of both types, especially common in CM.

Similarly, projection of the excitatory neuron PC1 from Fig. 1e onto the spatial transcriptomics data showed that the primary end of the gradient mapped to MD, especially MDmc, while the tertiary end mapped to PVT and PT (Supplementary fig. 2b). Among the surrounding nuclei, intralaminar nuclei were on the tertiary end, while motor/somatosensory and lateral dorsal thalamus were on the primary end, consistent with their profiles in mouse thalamus^8^. RNA-FISH confirmed this spatial distribution of marker genes associated with opposite ends of the gradient and showed that numerous cells co-expressed multiple markers (Fig. 2b, Supplementary fig. 3). These analyses suggest that the gene expression gradient of excitatory neurons corresponds to a spatial gradient of neurons between MD and PT/PVT.

**Fig. 3.**
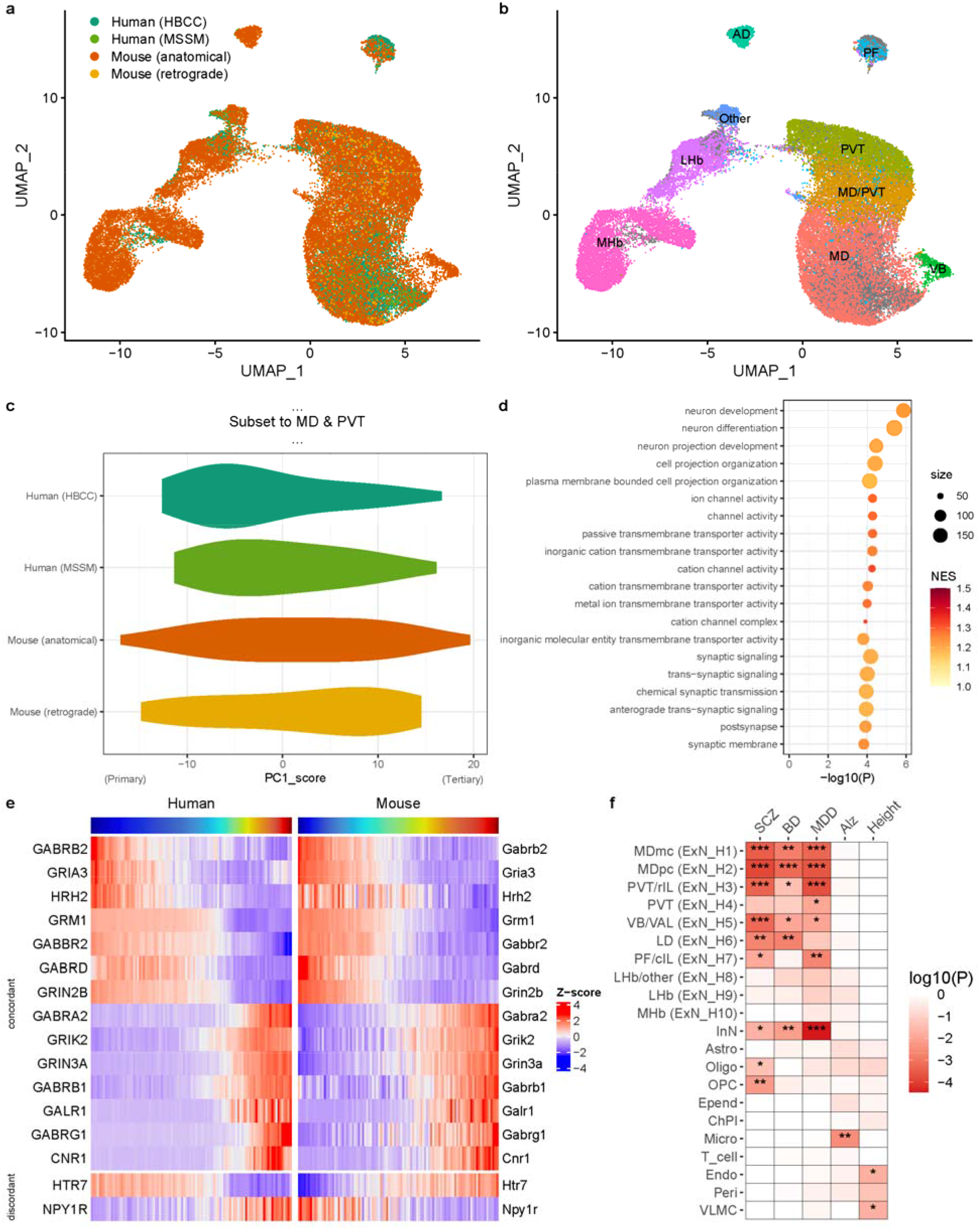
Comparison of mouse and human thalamic cell-type gradients and functional annotation. a. UMAP plot illustrating cross-species integration of human and mouse excitatory neurons in medial thalamus, colored by dataset: human data from HBCC and MSSM, and mouse data from this study based on anatomical dissections and a previous study using retrograde labeling from prefrontal cortex^8^. b. UMAP plot of integrated mouse and human data, with mouse cells labeled by their nuclei of origin inferred from integration with public datasets and Allen Brain Atlas ISH data (see also Supplementary fig. 4). Marker genes for individual mouse clusters are shown in Supplementary table 2. c. Violin density plot depicting distribution of PC1 scores for cross-species integrated dataset subset to excitatory neurons from MD or PVT across both human and mouse datasets. The ‘primary’ end is on the negative (left) and the ‘tertiary’ end on the positive (right) side of the x-axis. d. Gene ontology enrichment analysis showing the top 20 terms associated with PC1 from (c) using gene set enrichment analysis. Terms are ordered by theme (neuron projection development; ion channel activity; to synaptic function) and p-value. Size indicates number of genes per term, and color shows the normalized enrichment score (NES). The full test results are shown in supplementary table 3. e. Expression of genes along the cross-species PC1 between MD and PVT, as in (c) and (d), averaged across 100 bins of PC1 scores for each species is shown. Only neurotransmitter receptor-encoding genes with Spearman’s correlation >0.75 (concordant) or <0.5 (discordant) between human and mouse expression across bins are shown. f. MAGMA gene set test results show association of the top 10% most specific gene sets for each cell type with gene-level signal across five large GWAS (SCZ = schizophrenia; BD = bipolar disorder; MDD = major depressive disorder; Alz = Alzheimer’s disease). Asterisks denote significance levels (***: p<.001, **: p<.01, *: p<.05). The full test results are shown in Supplementary table 4.

Having identified a cell-type gradient in human MD and PVT with major similarities to the thalamic cell-type gradient in mice, we sought to better understand the commonalities and differences in MD cellular composition between humans and mice. Since previous snRNA-seq from retrogradely labeled cells did not include enough neurons in MD^8^, we collected additional snRNA-seq from mouse MD based on anatomical dissections. As expected, this identified the same major cell classes as human MD, except that inhibitory neurons were not observed in mouse MD (Supplementary fig. 4). To directly compare thalamic projection neurons between species, we integrated human and mouse excitatory neurons (Fig. 3a, Supplementary fig. 5) and assigned the resulting clusters to their putative nuclei of origin based on integration with existing data from mouse PVT^13^ and habenula^18^ and on marker gene expression in the Allen Institute’s ISH database. This allowed us to identify a correspondence between human and mouse thalamic neurons, confirming clusters derived from MD, intralaminar nuclei, and PVT, and demonstrating that additional discrete clusters corresponded to the parafascicular nucleus, lateral habenula, and medial habenula (Fig. 3b, Supplementary fig. 4c-f).

Cells within MD and PVT formed a gradient captured by PC1 with loadings closely associated with the primary-secondary-tertiary gradient described above (Supplementary fig. 5c-d). MD largely occupied the primary side, while PVT occupied the tertiary side of the gradient. Notably, the primary end of the gradient is expanded in humans relative to mice (Fig. 3c).

Gene ontology analysis showed that the genes associated with PC1 across both species were enriched for neuron projection development, ion channels, and synaptic transmission (Fig. 3d). Since many genes related to synaptic transmission were associated with this gradient, we compared neurotransmitter receptor gene expression between the species. Most genes showed a concordant pattern along the MD-PVT gradient between both species, but a few genes were discordant or species-specific in their gradient association (Fig. 3e, Supplementary fig. 5e).

Thus, while the gene expression patterns along the gradient are largely conserved, subtle differences in neuropeptide/neuromodulator receptors exist between species.

Finally, we examined which thalamic cell types may be most relevant to common genetic risk for serious mental illness. We used MAGMA^19^ to analyze gene-level summary statistics from the most recent genome-wide association studies (GWAS) of schizophrenia, bipolar disorder, and major depressive disorder. We tested for association with cell type specificity scores derived from our data using the top 10% most specific gene sets, as previously described^20^. Neuronal populations in the thalamus were significantly associated with genetic risk for severe mental illness: ExN_H2 (MDpc-enriched) was most strongly associated with genetic risk for schizophrenia and bipolar disorder, while thalamic inhibitory neurons and ExN_H3 (PVT-enriched) were most strongly associated with genetic risk for major depressive disorder. GWAS for Alzheimer’s disease and height, included for control purposes, implicated microglia and vascular cells, respectively, as expected.

We produced a cell type atlas of adult human MD and adjacent midline nuclei, spatially mapped the major neuronal populations, and compared mouse and human neuronal identities. Recent studies examined cell populations across the entire human brain^15^, fetal human thalamus^11^, and multiple structures within the mouse thalamus^8,13,14,21–23^ and habenula^18,24^, but were not focused on the MD, and cross-species comparisons have been limited to visual thalamus^21^, making our study the first in-depth analysis of cell types in human MD.

We identified a major transcriptional gradient between MD and PVT, which closely aligns with a previously-described gradient from primary to tertiary thalamus and likely reflects distinct connectivity and functional properties of these neurons^8^. This gene expression gradient mapped onto anatomical space, and respective clusters were enriched in MDmc, MDpc, and adjacent nuclei. Whilst MD has historically been considered ‘higher-order’, we identify major primary cell populations within both mouse and human MD. Cross-species analyses show that primary cells in MD are substantially expanded in humans relative to mice. This work has important functional implications as primary thalamic neurons tend to project densely to middle cortical layers and innervate fast-spiking interneurons, a property also observed for MD neurons^25,26^. Their expansion in human MD may thus be tied to granular prefrontal cortex expansion in anthropoid primates^27^.

Another obvious difference between species was the abundance of midbrain-derived inhibitory neurons in human MD, particularly MDpc and MDdc. Common genetic risk for schizophrenia and bipolar disorder was enriched in MD excitatory neurons, especially a cluster mapping onto MDpc. These results are in line with previous reports of neuronal loss in schizophrenia being more pronounced in MDpc than MDmc^28^. Genetic risk for major depressive disorder was strongly associated with PVT/PT neurons, consistent with their connectivity to limbic areas, and thalamic inhibitory interneurons, reinforcing similar findings in fetal human brain^29^ and suggesting the likely relevance of this species-divergent neuronal population linked to local thalamic circuit complexity^30^ for human psychopathology.

Limitations of this study include dissection variability across samples, modest sample size, and limited precision of gene-level GWAS risk calculation. Future work is needed to address these limitations, better understand the gene regulatory and neurodevelopmental basis of the observed cell-type gradient, and elucidate any changes in psychiatric illness.

## Methods

### Human brain tissue samples

Human brain samples were collected under protocols approved by the Institutional Review Boards (IRB) or the HBCC Oversight Committee with permission of the next-of-kin. Neurotypical controls were defined as having no history of psychiatric or neurological disorder and negative toxicology for psychotropic drugs. Demographics are summarized in Supplementary table 2.

### Mouse brain tissue samples

Experimental procedures were approved by the National Institute of Mental Health (NIMH) Animal Care and Use Committee. Mice were housed under a 12-hour light-dark cycle with food and water available ad libitum. Three-month-old wild-type C57BL/6NJ mice (The Jackson Laboratory) were used for all experiments.

### Dissection, nuclei isolation, and snRNAseq

For HBCC brain samples, the MD and adjacent midline nuclei were dissected from 1 cm thick coronal slabs as illustrated in Fig. 1a. Approximately 100mg of tissue (roughly 10-20% of the of the dissected volume) were used for nuclei isolation. To evenly represent the structure, the dissected tissue was crushed, and small tissue chunks were aliquoted randomly. For MSSM samples, pulverized brain tissue samples from medial thalamus were obtained through NeuroBioBank, and aliquots of 100mg tissue were used. Due to the high myelin content in human brain tissue, nuclei isolation involved a sucrose density gradient centrifugation using the Nuclei PURE nuclei isolation kit (Sigma-Aldrich #NUC201), as previously described^31^.

For mouse samples, MD and adjacent midline nuclei were dissected from 200 µm thick coronal sections from eight mice (four males, four females). To ensure sufficient nuclei yield, samples from the four males and four females, respectively, were pooled. Nuclei were isolated using a previously described protocol involving a less aggressive density gradient centrifugation^32^. Two samples from each pool of four animals were created.5,000-10,000 nuclei were loaded on the Chromium Controller (10X Genomics), and snRNA-seq libraries prepared using Chromium Next GEM Single Cell 3L Kits (v3.1 Chemistry Single Index). Libraries were sequenced on a NextSeq 2000 system (Illumina), followed by targeted deeper sequencing on a NovaSeq 6000 system (Illumina) for the ten human samples used.

### snRNAseq data quality control, and filtering

snRNA-seq data were mapped and gene-level unique molecular identifier (UMI) counts quantified using *CellRanger* (v6.1.1 and v7.1.0) and GRCh38 (GENCODE v32/Ensembl98) and mm10 (GENCODE vM23/Ensembl98) as reference genomes for human and mouse, respectively. Ambient RNA contamination was inferred and removed for each sample using *SoupX*^33^ with default settings. To ensure high-quality nuclei, an iterative filtering and clustering approach was used. First, nuclei were required to have > 500 UMI counts, >200 detected genes (nGenes), and < 5% rate of genes mapping to mitochondrial DNA (mtDNA rate). In the second step, doublets were predicted using *scDblFinder* (with doublet score thresholds determined separately for each sample with default settings) and removed along with clusters with mean number of detected genes <800 and mean mtDNA rate >2% (HBCC and mouse) or >2.5% (MSSM; due to higher mtDNA rate in those samples) were removed. In a third step, potential doublet clusters with markers of multiple cell classes (astrocytes fused with microglia, oligodendrocytes, or OPCs) were removed along with mtDNA-encoded and ribosomal subunit-encoding genes to ensure that clustering was not driven by these genes. At each step, *Harmony* integration^34^ of the top 2,000 most variable genes was used to account for individual sample-level batch effects, and shared nearest neighbor (SNN) clustering using the Louvain algorithm, a resolution of 0.8, and the top 30 dimensions, was applied to define clusters in *Seurat* (v4)^35^.

### Cell class annotation and MSSM sample inclusion

Major cell classes were assigned based on expression of canonical marker genes. Most samples contained excitatory neurons expressing *SLC17A6* (VGluT2) and inhibitory neurons expressing *SOX14*. However, two MSSM samples (MSSM3 and MSSM4) contained mostly VGluT1-positive excitatory neurons and *SOX14*-negative inhibitory neurons (Supplementary fig. 1a-b). Integration with Human Brain Cell Atlas data from Siletti et al. (2023)^15^ showed that these cells may stem from telencephalic structures (Supplementary fig. 1c-d). Samples MSSM3 and MSSM4 were therefore excluded from downstream analyses.

### Integration between datasets

#### Human Brain Cell Atlas integration

Data from the medial thalamus and epithalamus regions of interest of the Human Brain Cell Atlas^15^ were downloaded and integrated with the HBCC and MSSM data, respectively, using anchors defined via canonical correlation analysis (CCA) in *Seurat* (Supplementary fig. 1c). Nearest centroid classification of HBCC and MSSM cells by supercluster label was performed to map corresponding cell types (Supplementary fig. 1d).

#### Brain bank integration

Excitatory neurons from HBCC and MSSM were integrated using CCA-based anchors in *Seurat*. SNN clustering with a resolution of 0.5 on the top 5 PCs was used to define the clusters. To alleviate clustering being driven by individual samples, the four most closely related clusters were merged, resulting in 10 clusters (Supplementary fig. 1e-f).

#### Mouse anatomical and retrograde labeling data integration

Mouse thalamic neurons obtained in this study were integrated with those based on retrograde labeling from prefrontal cortex in a previous study^8^ using CCA-based anchors in *Seurat*. SNN clustering with a resolution of 0.8 on the top 10 PCs was used, resulting in 19 clusters (Supplementary fig. 4b).

#### PVT data integration

Five populations previously annotated and validated in the mouse PVT^13^ were integrated with the mouse thalamic neuron clusters using anchors based on reciprocal principal component analysis (RPCA) in *Seurat*. Nearest centroid classification was used to map the PVT clusters onto corresponding excitatory neuron clusters (Supplementary fig. 4c).

#### Habenula data integration

Previously annotated and validated neuron populations from the mouse habenula^18^ were re-identified and integrated with the mouse thalamic neurons using RPCA-based anchors. Nearest centroid classification was used to map the re-identified habenula populations onto the excitatory neuron clusters (Supplementary fig. 4d).

All integrations were performed on the top 2,000 most variable genes across datasets (SelectIntegrationFeatures in *Seurat*). Nearest neighbor classifiers were implemented via the *lolR* library (lol.classify.nearestCentroid) on the top PCs of the integrated data (top 30 PCs for cell class labels, and top 10 PCs for excitatory neuron labels).

### Cross-species analyses

Cross-species integration of human and mouse excitatory neurons was achieved by first restricting datasets in both species to high-confidence one-to-one orthologs based on their BioMart annotation (Ensembl105), followed by merging and integration of the datasets using CCA-based anchors in *Seurat* (Fig. 3a; Supplementary fig. 5a). Nearest neighbor classification was used to map each mouse excitatory neuron cluster (or prefrontal-projecting cluster^8^) onto the human excitatory neuron clusters (Supplementary fig. 5b). The same type of integration with CCA-based anchors was then applied again, but only to the clusters inferred to be derived from MD and PVT (ExN_H1-4, ExN_M1-7; Fig. 3c, Supplementary fig. 5c).

Cross-species comparison of PC1 gene loadings (Supplementary figs. 1h and 5d) were performed on the intersection of the genes across which the PCs were calculated (top 2,000 variable genes and the published list of top 500 differentially expressed genes across mouse thalamic nuclei^8^) and restricted to high-confidence one-to-one orthologous genes.

### Marker gene testing

Marker genes for each cell type label were obtained in *Seurat* using a Wilcoxon rank sum test. Only positive markers with a minimum log fold change of 0.25 and detected in at least 25% of the respective cell population were included (Supplementary data tables 1 and 2).

### Functional gene set enrichment

Gene set enrichment analysis of PC1-associated genes was performed using the *fgsea* R package^36^ with human gene ontology annotations obtained through *org*.*Hs*.*eg*.*db* (v3.11.4), absolute PC1 loading values as the ranking metric, and the following options: minSize=10, maxSize=200, scoreType=“pos”. Test statistics are shown in Supplementary table 3.

### MAGMA gene set tests

To perform genetic associations of cell type-specific gene sets with gene-level signal from GWAS, summary statistics for schizophrenia^37^, bipolar disorder^38^, major depressive disorder^39^ (without 23andMe data), Alzheimer’s disease^40^, and height^41^ were downloaded and reformatted using the *MungeSumstats*^42^ package. The *MAGMA*.*Celltyping*^20^ package was used to obtain reference genome plink files (based on the 1000 Genomes European population), and call *MAGMA*^19^ (v1.10) to link SNPs to genes and obtain gene level trait associations. Cell type specificity scores were then calculated using the *EWCE*^43^ package and used for top 10% gene set tests for each cell type via *MAGMA*.*Celltyping*. A window size of 35kb upstream and 10kb downstream was used for gene definitions. Test results are shown in Supplementary table 4.

### Cryo-sectioning and histology

Blocks of thalamus tissue were sectioned on a cryostat. 14µm thick frozen coronal sections were generated at multiple anterior-posterior levels. A combined myelin and neuronal cell body stain (luxol-fast blue and cresyl violet^44^) were utilized to determine their anterior-posterior position of adjacent sections based on a neuroanatomical atlas of the human brain^45^ and select two sections containing the MD for multiplexed RNA-FISH and spatial transcriptomics.

Brightfield histological stains were imaged on a Zeiss Axioscan Z1 widefield slide scanner system (software: ZEN v3.1) and implementing autofocus, online stitching and shading correction. All images were acquired using a 10X objective and collected as 16-bit.

### Spatial transcriptomics data acquisition

Two 14µm thick frozen coronal sections were fixed in 100% methanol at -20□ C for 30 minutes and a hematoxylin and eosin (H&E) staining was applied following the Visium CytAssist Spatial Gene Expression for Fresh Frozen Protocol (CG000614, 10X Genomics). Samples were then processed via the Visium CytAssist Spatial Gene Expression Reagent Kit for Human Transciptome with an 11×11mm wide capture area (CG000495, 10X Genomics). Libraries were sequenced on a NextSeq 2000 system (Illumina).

### Spatial transcriptomics data analysis

Visium data were processed using *SpaceRanger* (v2.1.1) to map reads to the human probe set (v2.0), generate gene-level UMI counts per spot, and align the fiducial frame to locate the spatial position of the barcodes on the H&E stain image. To ensure high-quality data, only spots with a >5,000 UMI counts, >2,500 detected genes, and <15% mtDNA rate were kept. Spots were then subjected to robust cell type decomposition (RCTD)^17^ via the *spacexr* package, which uses non-negative least squares regression for deconvolution, using our snRNA-seq data from human MD with annotated cell type labels as the reference dataset. The spatial distribution of predicted cell type proportions (RCTD weights) is shown in Fig. 2a and Supplementary fig. 3a. Data was then normalized using *sctransform*^46^. Normalized gene expression values per spot were used for visualization of individual gene expression levels and projection onto PC1 from the excitatory neuron snRNA-seq data (Supplementary fig. 2b).

### Multiplexed RNA-FISH data acquisition

Two 14µm thick frozen coronal sections were post-fixed in 4% paraformaldehyde and processed for multiplexed RNA-FISH using the RNAscope HiPlex v1 kit (Advanced Cell Diagnostics), as previously described^13^. Fluorescently labeled probes targeting mRNA of *GRM1* (AlexaFluor488), *SNCA* (ATTO550), *SHISA6* (ATTO647), and *FOXP2* (AlexaFluor750) along with a nuclear stain (DAPI) were imaged on a Nikon Biopipeline Slide system at 40X resolution. Additionally, three ‘nuisance channels’ were used to capture autofluorescence. The BioPipeline was operated via Nikon Elements with a custom-programmed automated acquisition setting using specified shading correction, exposure times, autofocusing, z-stack settings, and regions of interest. The 4X objective was utilized to conduct a low-resolution scan, and the image was used to delineate the regions of interest within the sample. All acquisitions were performed using the 20X objective. LED power levels and camera exposure times were adjusted to minimize saturation in areas of the brightest signal on the section, and shading corrections were individually created for each channel. Images were acquired in 16-bit format, with a scaled pixel size of 0.32μm. Eight planes were acquired with a Z-step size of 2μm.

### Multiplexed RNA-FISH data analysis

The acquired .nd2 files were stitched using the “blend” algorithm in NIS Elements and corrected for extended depth of field. The samples showed a widespread occurrence of highly autofluorescent signal, presumably from lipofuscin accumulation. We computed multiple masks for the autofluorescence based on the three acquired ‘nuisance channels’ by thresholding these channels at various percentiles. Appropriate thresholds were then chosen by visual inspection (AS). The corresponding masks of the autofluorescence were used to remove it from all other channels. All pixels with identified non-specific intensities were masked to create an intensity value of 0. Multichannel .tiff files were then imported into Arivis Vision4D software (v4.1.1).

Using the Machine Learning Trainer module, an experimenter (SKWA) created training data for random forest model creation for each channel by manually identifying clouds of mRNA puncta over and surrounding DAPI-labeled nuclei as transcript-positive cells. Each channel’s model was applied to the appropriate image to identify transcript-positive cells. The overlap of the ‘cells’ was identified by the percent overlap of the segmentations (> 50%). Lookup tables and gamma were adjusted for contrast in the creation of Fig. 2b and Supplemental fig. 3.

## Data and code availability

De-identified human single-nucleus and spatial sequencing data will be deposited on NIMH Data Archive (NDA). Mouse single-nucleus data will be deposited on the Gene Expression Omnibus (GEO), and analysis code will be made available on GitHub at the time of publication.

## Contributions

AS initiated and designed the study, assisted with data acquisition, analyzed data, and wrote the paper. AS, PKA, SM, and FJM designed the study. NF, PKA, YL, QX, and VI acquired human single-nucleus and/or spatial data. NF, AE, JS, and MCK prepared single-nucleus and spatial libraries. AM and RK acquired mouse single-nucleus data. SKWA, SR, and TBU acquired and analyzed imaging data. PKA, AM, RK, CG, YP, NA, AR, VM, PR, LD, MMH, SH, MAP, SM, and FJM assisted with data analysis and interpretation. PKA, MMH, MAP, SM, and FJM edited the paper.

## Acknowledgements

We thank the families who donated the brain of their loved ones for research, Richard Edelmann and Jonathan Kuo for assistance with imaging data acquisition, Bayu Sisay for assistance with library preparation, Mark Cookson, Rebekah Langston, Ariel Levine, and Kaya Matson for help with optimization of nuclei isolation protocols, Siyuan Liu, Alex DeCasien, Aleksandra Dakic, Gabi Dugan, Pranav Narnur, Rachel Smith, Bruno Averbeck, Mary Baldwin, James Bourne, and all members of the Human Genetics Branch and HBCC for feedback on the project, Ajeet Mandal, Barbara Lipska, Vahram Haroutunian and the NeuroBioBank team for brain banking resources, and the NIH HPC Biowulf team for computational resources. This research was supported by the Intramural Research Program of the NIMH (ZIA-MH002810 [FJM], ZIC-MH002903 [SM], and ZIA-MH002950 [MAP]), FNLCR Contract 75N91019D00024 (MCK), and the following extramural grants: K22-MH126015 (AS), K99-MH129613 (AM), R01-MH120118 (MMH), and P50-MH132642 (MMH).

## Supplementary figure captions

- Supplementary fig. 1 – Human MD and midline thalamus cell type identification and comparison to existing datasets.
- Supplementary fig. 2 – Mapping of neuronal populations and gene expression gradients using spatial transcriptomics.
- Supplementary fig. 3 – Analysis of co-expression in multiplexed fluorescent in situ hybridization data.
- Supplementary fig. 4 – Mouse MD and midline thalamus cell type identification and comparison to existing datasets.
- Supplementary fig. 5 – Cross-species integration of mouse and human cell types in MD and midline thalamus.

## Supplementary table captions

- Supplementary table 1 – Summary of major spatial mapping results for MD and adjacent thalamic nuclei.
- Supplementary table 2 – Demographics summary of human brain samples.

## Supplementary data table captions

- Supplementary data table 1 – Marker genes for human MD clusters
- Supplementary data table 2 – Marker genes for mouse MD clusters
- Supplementary data table 3 – Gene ontology enrichment analysis
- Supplementary data table 4 – MAGMA gene set test statistics

## References

1. Parnaudeau, S., Bolkan, S. S. & Kellendonk, C. The Mediodorsal Thalamus: An Essential Partner of the Prefrontal Cortex for Cognition. Biol. Psychiatry 83, 648–656 (2018).

2. Wolff, M. & Halassa, M. M. The mediodorsal thalamus in executive control. Neuron 112, 893–908 (2024).

3. Penzo, M. A. & Gao, C. The paraventricular nucleus of the thalamus: an integrative node underlying homeostatic behavior. Trends Neurosci. 44, 538–549 (2021).

4. Huang, A. S. et al. Thalamic Nuclei Volumes in Psychotic Disorders and in Youths With Psychosis Spectrum Symptoms. Am. J. Psychiatry 177, 1159–1167 (2020).

5. Woodward, N. D. & Heckers, S. Mapping Thalamocortical Functional Connectivity in Chronic and Early Stages of Psychotic Disorders. Biol. Psychiatry 79, 1016–1025 (2016).

6. Anticevic, A. & Halassa, M. M. The thalamus in psychosis spectrum disorder. Front. Neurosci. 17, (2023).

7. Crail-Melendez, D., Atriano-Mendieta, C., Carrillo-Meza, R. & Ramirez-Bermudez, J. Schizophrenia-like psychosis associated with right lacunar thalamic infarct. Neurocase 19, 22–26 (2013).

8. Phillips, J. W. et al. A repeated molecular architecture across thalamic pathways. Nat. Neurosci. 22, 1925–1935 (2019).

9. Oldham, S. & Ball, G. A phylogenetically-conserved axis of thalamocortical connectivity in the human brain. Nat. Commun. 14, 6032 (2023).

10. Govek, K. W. et al. Developmental trajectories of thalamic progenitors revealed by single-cell transcriptome profiling and Shh perturbation. Cell Rep. 41, 111768 (2022).

11. Kim, C. N., Shin, D., Wang, A. & Nowakowski, T. J. Spatiotemporal molecular dynamics of the developing human thalamus. Science 382, eadf9941 (2023).

12. Mukherjee, A., Lam, N. H., Wimmer, R. D. & Halassa, M. M. Thalamic circuits for independent control of prefrontal signal and noise. Nature 600, 100–104 (2021).

13. Gao, C. et al. Molecular and spatial profiling of the paraventricular nucleus of the thalamus. eLife 12, e81818 (2023).

14. Shima, Y. et al. Distinctiveness and continuity in transcriptome and connectivity in the anterior-posterior axis of the paraventricular nucleus of the thalamus. Cell Rep. 42, 113309 (2023).

15. Siletti, K. et al. Transcriptomic diversity of cell types across the adult human brain. Science 382, eadd7046 (2023).

16. Goldman-Rakic, P. S. & Porrino, L. J. The primate mediodorsal (MD) nucleus and its projection to the frontal lobe. J. Comp. Neurol. 242, 535–560 (1985).

17. Cable, D. M. et al. Robust decomposition of cell type mixtures in spatial transcriptomics. Nat. Biotechnol. 40, 517–526 (2022).

18. Wallace, M. L. et al. Anatomical and single-cell transcriptional profiling of the murine habenular complex. eLife 9, e51271 (2020).

19. Leeuw, C. A. de, Mooij, J. M., Heskes, T. & Posthuma, D. MAGMA: Generalized Gene-Set Analysis of GWAS Data. PLOS Comput. Biol. 11, e1004219 (2015).

20. Skene, N. G. et al. Genetic identification of brain cell types underlying schizophrenia. Nat. Genet. 50, 825–833 (2018).

21. Bakken, T. E. et al. Single-cell and single-nucleus RNA-seq uncovers shared and distinct axes of variation in dorsal LGN neurons in mice, non-human primates, and humans. eLife 10, e64875 (2021).

22. Kapustina, M. et al. The cell-type-specific spatial organization of the anterior thalamic nuclei of the mouse brain. Cell Rep. 43, (2024).

23. Mandelbaum, G. et al. Distinct Cortical-Thalamic-Striatal Circuits through the Parafascicular Nucleus. Neuron 102, 636-652.e7 (2019).

24. Hashikawa, Y. et al. Transcriptional and Spatial Resolution of Cell Types in the Mammalian Habenula. Neuron 106, 743-758.e5 (2020).

25. Delevich, K., Tucciarone, J., Huang, Z. J. & Li, B. The mediodorsal thalamus drives feedforward inhibition in the anterior cingulate cortex via parvalbumin interneurons. J. Neurosci. Off. J. Soc. Neurosci. 35, 5743–5753 (2015).

26. Collins, D. P., Anastasiades, P. G., Marlin, J. J. & Carter, A. G. Reciprocal Circuits Linking the Prefrontal Cortex with Dorsal and Ventral Thalamic Nuclei. Neuron 98, 366-379.e4 (2018).

27. Preuss, T. M. & Wise, S. P. Evolution of prefrontal cortex. Neuropsychopharmacology 47, 3–19 (2022).

28. Popken, G. J., Bunney, W. E., Potkin, S. G. & Jones, E. G. Subnucleus-specific loss of neurons in medial thalamus of schizophrenics. Proc. Natl. Acad. Sci. 97, 9276–9280 (2000).

29. Mannens, C. C. A. et al. Chromatin accessibility during human first-trimester neurodevelopment. Nature (2024) doi:10.1038/s41586-024-07234-1.

30. Arcelli, P., Frassoni, C., Regondi, M. C., Biasi, S. D. & Spreafico, R. GABAergic Neurons in Mammalian Thalamus: A Marker of Thalamic Complexity? Brain Res. Bull. 42, 27–37 (1997).

31. Langston, R. G. et al. Association of a common genetic variant with Parkinson’s disease is mediated by microglia. Sci. Transl. Med. 14, eabp8869 (2022).

32. Matson, K. J. E. et al. Isolation of Adult Spinal Cord Nuclei for Massively Parallel Single-nucleus RNA Sequencing. J. Vis. Exp. JoVE 58413 (2018) doi:10.3791/58413.

33. Young, M. D. & Behjati, S. SoupX removes ambient RNA contamination from droplet-based single-cell RNA sequencing data. GigaScience 9, giaa151 (2020).

34. Korsunsky, I. et al. Fast, sensitive and accurate integration of single-cell data with Harmony. Nat. Methods 16, 1289–1296 (2019).

35. Hao, Y. et al. Integrated analysis of multimodal single-cell data. Cell 184, 3573-3587.e29 (2021).

36. Korotkevich, G. et al. Fast gene set enrichment analysis. 060012 Preprint at 10.1101/060012 (2021).

37. Trubetskoy, V. et al. Mapping genomic loci implicates genes and synaptic biology in schizophrenia. Nature 604, 502–508 (2022).

38. Mullins, N. et al. Genome-wide association study of more than 40,000 bipolar disorder cases provides new insights into the underlying biology. Nat. Genet. 53, 817–829 (2021).

39. Howard, D. M. et al. Genome-wide meta-analysis of depression identifies 102 independent variants and highlights the importance of the prefrontal brain regions. Nat. Neurosci. 22, 343–352 (2019).

40. Wightman, D. P. et al. A genome-wide association study with 1,126,563 individuals identifies new risk loci for Alzheimer’s disease. Nat. Genet. 53, 1276–1282 (2021).

41. Yengo, L. et al. A saturated map of common genetic variants associated with human height. Nature 610, 704–712 (2022).

42. Murphy, A. E., Schilder, B. M. & Skene, N. G. MungeSumstats: a Bioconductor package for the standardization and quality control of many GWAS summary statistics. Bioinformatics 37, 4593–4596 (2021).

43. Skene, N. G. & Grant, S. G. N. Identification of Vulnerable Cell Types in Major Brain Disorders Using Single Cell Transcriptomes and Expression Weighted Cell Type Enrichment. Front. Neurosci. 10, (2016).

44. Geisler, S., Heilmann, H. & Veh, R. W. An optimized method for simultaneous demonstration of neurons and myelinated fiber tracts for delineation of individual trunco- and palliothalamic nuclei in the mammalian brain. Histochem. Cell Biol. 117, 69–79 (2002).

45. Mai & Majtanik & Paxinos. Atlas Of The Human Brain, 4Th Edition. (Elsevier, 2016).

46. Hafemeister, C. & Satija, R. Normalization and variance stabilization of single-cell RNA-seq data using regularized negative binomial regression. Genome Biol. 20, 296 (2019).

