## Supplementary Figures and Tables for "A conserved cell-type gradient across the human mediodorsal and paraventricular thalamus"

#### Supplementary fig. 1

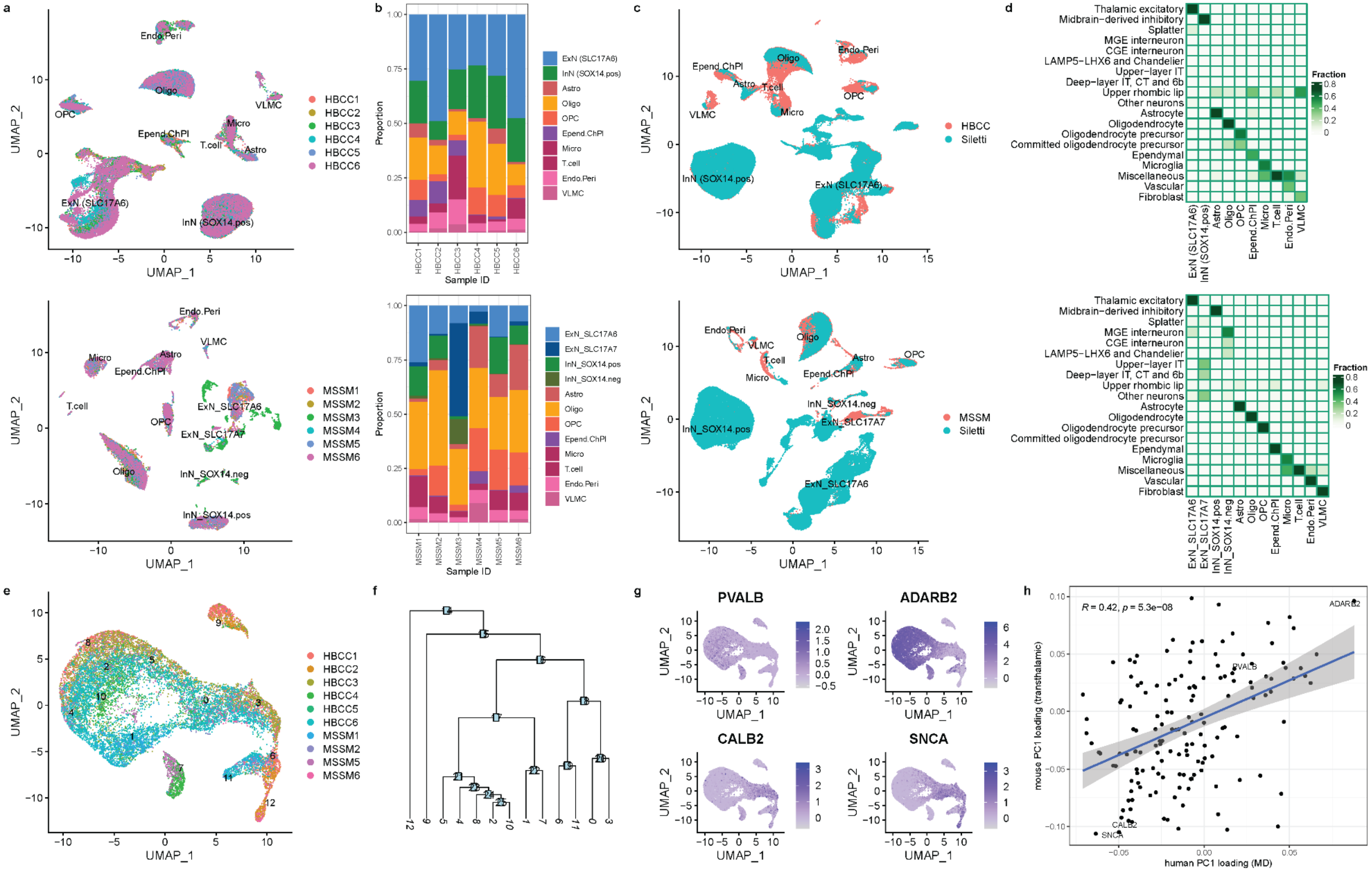

**Supplementary fig. 1 – Human MDT and midline thalamus cell type identification and comparison to existing datasets.**

a. UMAP plot of all cells that passed filtering and quality control (see Methods) labeled by annotated cell type and colored by individual sample from HBCC (top) and MSSM (bottom).
b. Distribution of cell type composition of each individual sample from HBCC (top) and MSSM (bottom).
c. UMAP plot illustrating the integration of samples from HBCC (top) and MSSM (bottom) with Human Brain Cell Atlas data from Siletti et al. (2023)[1] for medial thalamus and epithalamus, colored by annotated cell type.
d. Classification of cells from HBCC (top) and MSSM (bottom) based on superclusters from the Human Brain Cell Atlas data from the integrated data shown in (c). Samples MSSM3 and MSSM4 showed atypical excitatory and inhibitory cell types likely derived from outside the thalamus and virtually absent in all other samples. They were therefore removed from downstream analyses.
e. UMAP plot of excitatory neurons, colored by individual sample, and labeled by Seurat cluster number, indicating biases in cluster representation across individuals.
f. Dendrogram illustrating relationship between Seurat clusters shown in (e). The most closely related four clusters (2,4,8,10) were merged yielding the clusters shown in Fig. 1d.
g. UMAP plots of excitatory neurons, colored by normalized expression level of four genes previously associated with primary (*PVALB*, *ADARB2*) and tertiary (*CALB2*, *SNCA*) thalamic identities.
h. Scatter plot depicting correlation of human PC1 loadings (from excitatory neurons in MDT and adjacent nuclei) with mouse PC1 loadings (previously described thalamus-wide axis[2]) for the intersection of homologous variable genes across which PC1 was calculated in both datasets. A linear regression curve is shown with shading indicating 95% confidence interval of the slope. A Pearson's correlation coefficient (R) and significance in a two-sided t-test (p) are shown.

#### Supplementary fig. 2

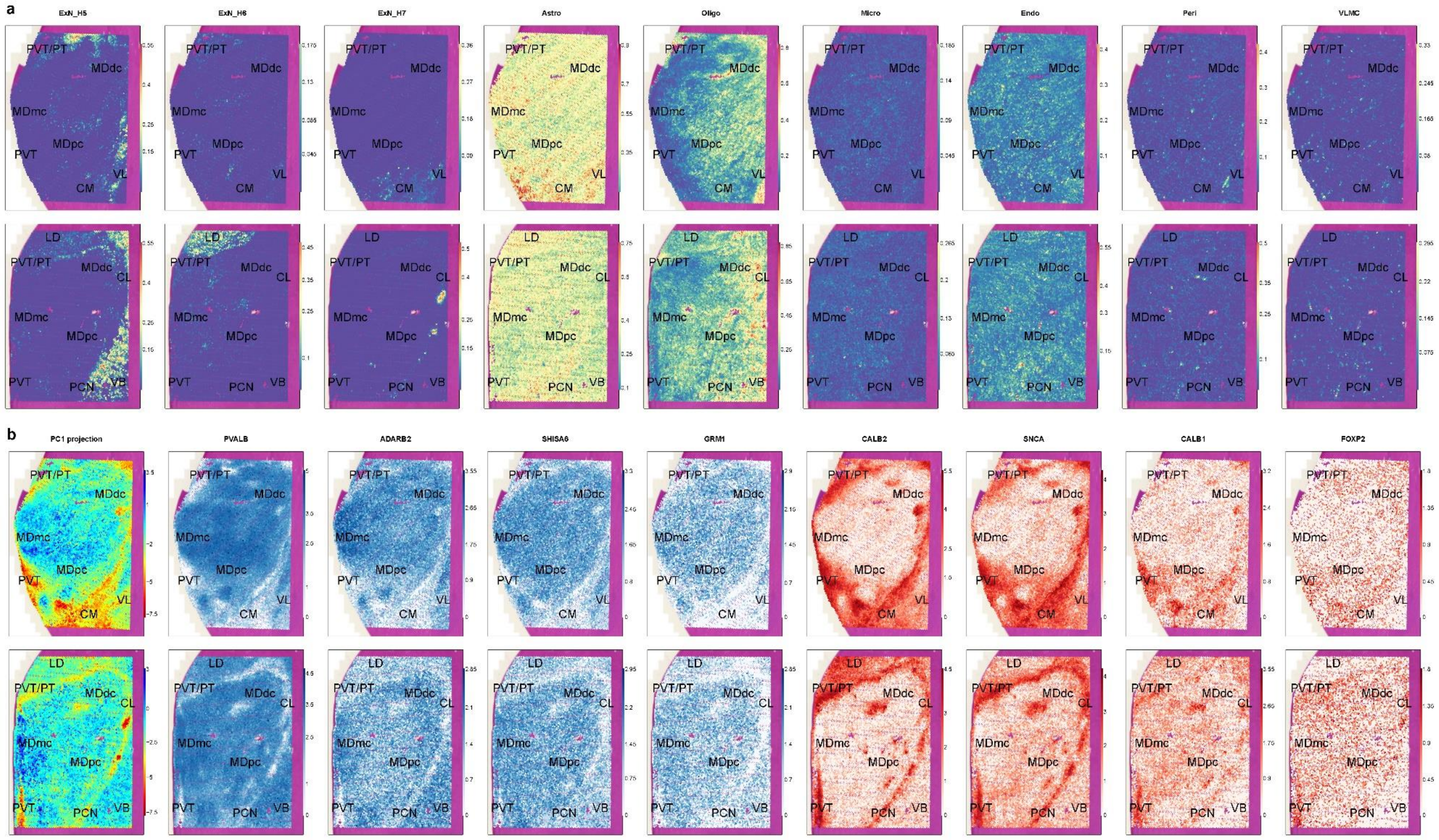

**Supplementary fig. 2 – Mapping of neuronal populations and gene expression gradients using spatial transcriptomics.**

a. Spatial mapping of inferred cell type proportions for three additional excitatory neuron populations (left three panels) and six major glial populations (right six panels) across two coronal sections (anterior section: top; posterior section: bottom). The clusters shown preferentially map onto motor/somatosensory nuclei (VL/VB, ExN_H5), lateral dorsal nucleus (LD, ExN_H6), and caudal subset of the intralaminar nuclei (ExN_H7)
b. Spatial mapping of projected PC1 scores form the snRNA-seq data in Fig. 1e-f and Supplementary fig. 1g-h (leftmost panel) and normalized expression level of four marker genes for primary thalamus (intensity colored in blue) and four marker genes for secondary/tertiary thalamus (intensity colored in red).

**Supplementary fig. 3**

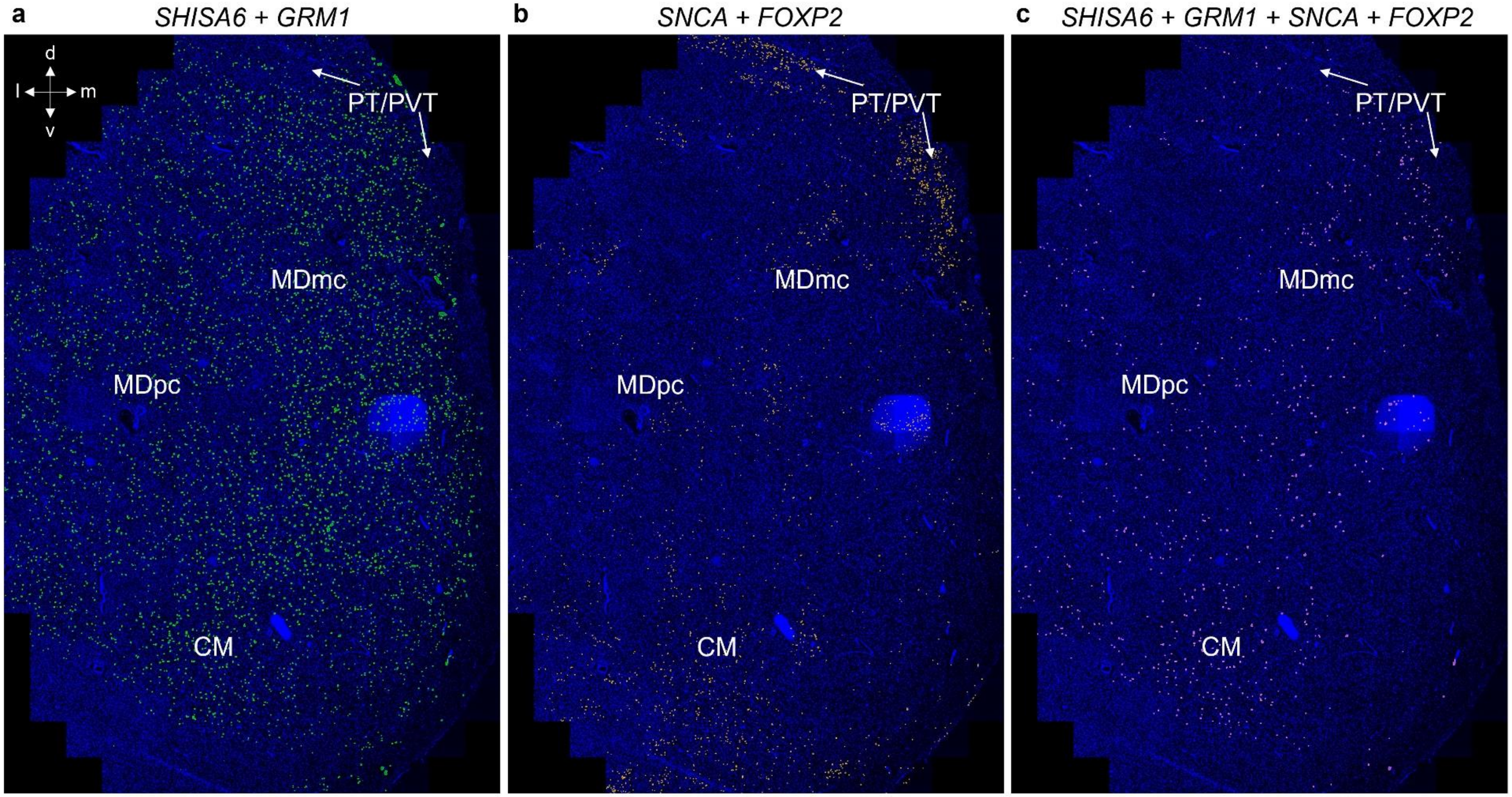

**Supplementary fig. 3 – Analysis of co-expression in multiplexed fluorescent in situ hybridization data.**

a. Co-expression of the two primary thalamus marker genes (*SHISA6* and *GRM1*), predominantly expressed in MDT, particularly in MDmc.
b. Co-expression of the two secondary/tertiary thalamus marker genes (*SNCA* and *FOXP2*), predominantly expressed in midline thalamus (PVT and PT), and intralaminar nuclei (CM).
c. Quadruple co-expression of both primary and tertiary markers in the same cells in the intralaminar nuclei and in the transition zone of MDT and PT/PVT.

### Supplementary fig. 4

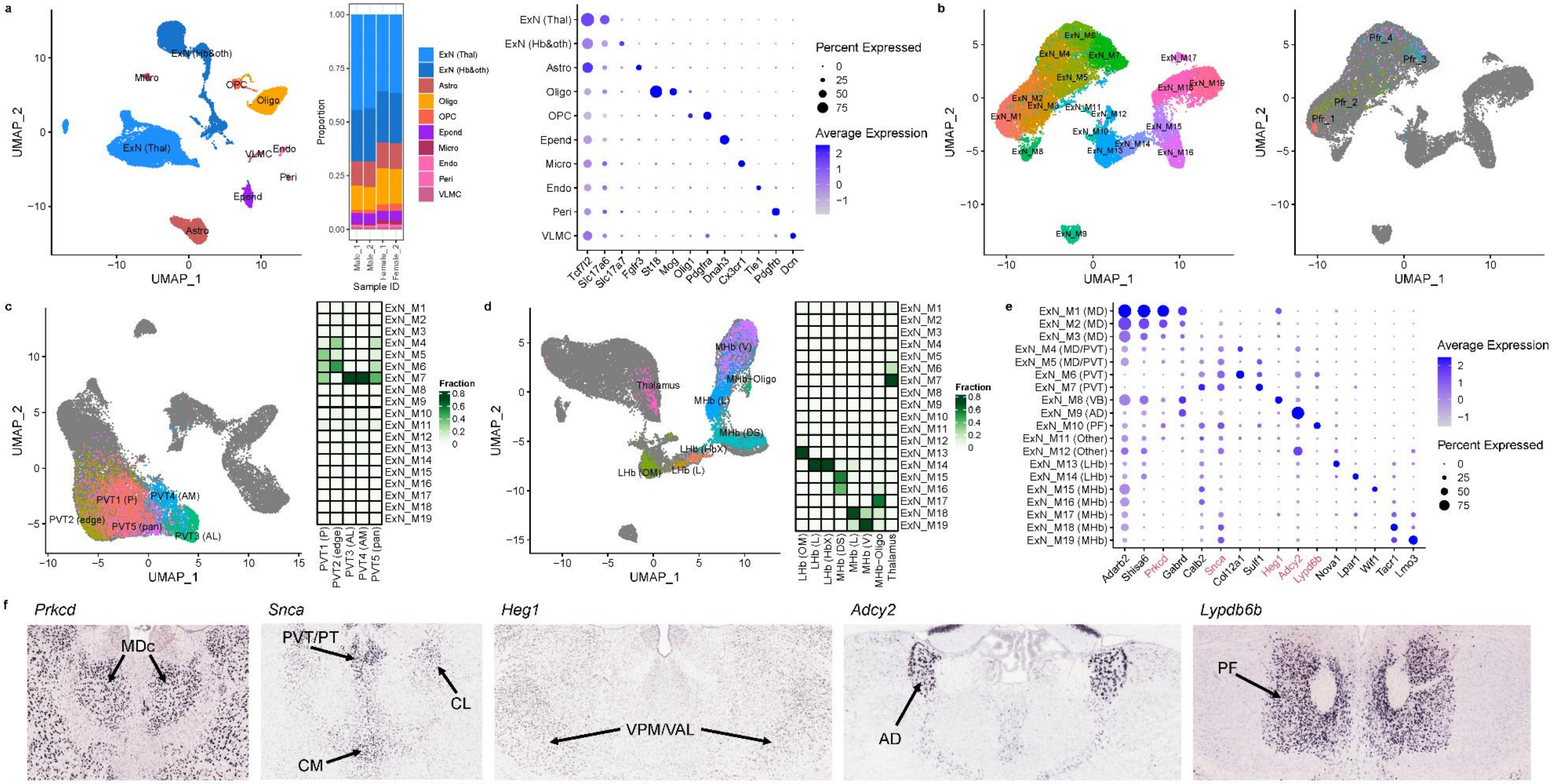

**Supplementary fig. 4 – Mouse MDT and midline thalamus cell type identification and comparison to existing datasets.**

a. UMAP plot of major cell classes in mouse MDT and midline thalamus (left), distribution of cell types across the four samples (middle), and dot plot showing expression levels of canonical marker genes for the major cell classes (right).
b. UMAP plots of excitatory neurons in mouse MDT and midline thalamus, colored by cluster identity (left) and previously defined cluster identifiers from retrogradely labeled cells from prefrontal cortex.
c. Integration of mouse MDT and thalamic midline excitatory neurons with existing data from the mouse PVT[3]. A UMAP plot of the integrated data (left) and classification based on mouse neuronal clusters (right) is shown. The five major PVT populations were: PVT1 (posterior), PVT2 (posteromedial edge), PVT3 (anterolateral), PVT4 (anteromedial), and PVT5 (pan-PVT).
d. Integration of mouse MDT and thalamic midline excitatory neurons with existing data from the mouse habenula[4]. Same plots as in (c) are shown. The lateral habenula (LHb) neurons included cells located in the oval/medial and marginal (OM), lateral (L), and borderline (HbX) portions. The medial habenula (MHb) neurons included cells in the dorsal/superior (DS), lateral (L), ventral (V) portions. Other populations stemmed from likely oligodendrocyte-contaminated cells and surrounding neurons from the thalamus.
e. Dot plot showing the expression of marker genes for excitatory neuron clusters in mouse MDT and midline nuclei. The nuclei origin of each cluster was inferred based on integration with PVT (c), habenula (d), and Allen Institute ISH data (f).
f. Allen Institute ISH data for five example marker genes. *Prkcd* is a marker for primary thalamus and labels cells within central MDT (MDc). *Snca* is a tertiary thalamus marker expressed in PVT/PT, and intralaminar nuclei (CM, CL, PCN). *Heg1* is a primary thalamus marker that is more highly expressed in primary nuclei outside of MDT (e.g., VPM). *Adcy2* is a marker for the anterior dorsal nucleus (AD), and *Lypdb6b* is a marker for the parafascicular nucleus.

### Supplementary fig. 5

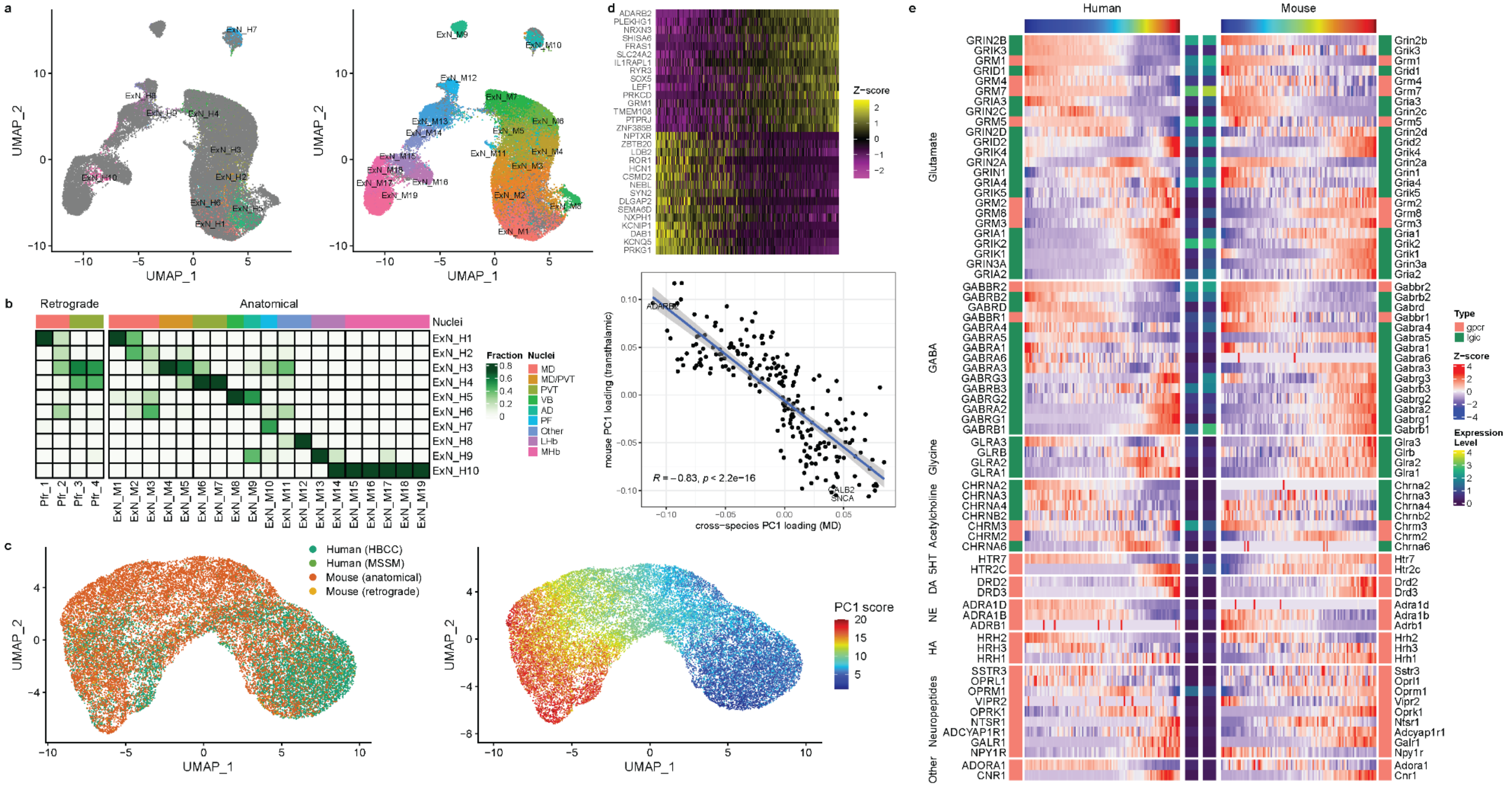

**Supplementary fig. 5 – Cross-species integration of mouse and human cell types in MDT and midline thalamus.**

a. UMAP plots, as in Fig. 3a-b, of integrated data from human and mouse excitatory neurons in medial thalamus, colored by human (left) and mouse (right) cluster identities.
b. Classification of mouse excitatory neuron populations from previously published retrogradely labeled cells from the prefrontal cortex[2] and anatomical dissections, based on human excitatory neuron populations.
c. UMAP plots of integrated data from human and mouse excitatory neurons subset to those inferred to be derived from MDT/PVT (ExN_H1-4 for humans, ExN_H1-7 for mice), colored by dataset and cross-species PC1 score. A density plot of human and mouse cells along PC1 is shown in Fig. 3c, demonstrating expansion of primary thalamic neurons in humans relative to mice.
d. Top: Heatmap of top 15 genes with highest and lowest loadings on the cross-species PC1, with cells are ordered by their PC1 score. Bottom: Scatter plot representing correlation of the cross-species PC1 loadings (cells from MDT and PVT) with previously published thalamus-wide PC1 loadings in mice[2]. A linear regression curve is shown with shading indicating 95% confidence interval of the slope. A Pearson's correlation coefficient (R) and significance in a two-sided t-test (p) are displayed.
e. Expression of genes along the cross-species PC1 between MDT and PVT, averaged across 100 bins of PC1 scores for each species, is shown. Row annotations denote receptor type (gpcr = g-protein coupled receptor; lgic = ligand-gated ion channel) and normalized expression level across all bins in human (center left) and mouse (center right). Only neurotransmitter receptor-encoding genes with an expression level above 0.05 in at least one species are shown. Rows are split based on receptor ligands (5HT = serotonin; DA = dopamine; NE = norepinephrine; HA = histamine).

**Supplementary Tables**

| Structure | Major projection target | InN density | ExN subcluster | Profile along gradient |
|---|---|---|---|---|
| **MDmc** | orbitofrontal, ventromedial PFC | | ExN_H1 | Primary end |
| **MDpc** | dorsolateral PFC | High | ExN_H2 | Primary |
| **MDdc/mf** | lateral-posterior PFC | High | ExN_H1 | Primary |
| **PVT/PT** | ventral striatum, amygdala, limbic cortex | | ExN_H3, ExN_H4 | Tertiary end |
| **VL/VB** | motor/somatosensory cortex | | ExN_H5 | Primary |
| **LD** | cingulate/retrosplenial cortex | | ExN_H6 | Primary |
| **rIL** | dorsal striatum, broad cortical | | ExN_H3 | Tertiary |
| **cIL** | dorsal striatum, broad cortical | | ExN_H7 | Tertiary |

**Supplementary table 1 – Summary of major spatial mapping results for MD and adjacent thalamic nuclei.**
MD subdivisions: mc = magnocellular, pc = parvocellular, dc/mf = densocellular/multiform; Other nuclei: PVT/PT = paraventricular/parataenial, LD = lateral dorsal, rIL = rostral intralaminar, cIL = caudal intralaminar.

| Dataset | Sex (f;m) | Ethnicity (Afr;Eur) | Age (mean ± SD) | PMI (mean ± SD) |
|---|---|---|---|---|
| **HBCC** | 4;2 | 4;2 | 35.7 ± 10.6 | 33.4 ± 10.4 |
| **MSSM** | 2;4 | 2;4 | 50.5 ± 12.7 | 12.7 ± 5 |

**Supplementary table 2 – Demographics summary of human brain samples.**
Acronyms: f = female, m = male, Afr = Black or African American, Eur = White or European, SD = standard deviation, Age = age of death in years, PMI = post-mortem interval in hours.

References:
1. Siletti, K. *et al.* Transcriptomic diversity of cell types across the adult human brain. *Science* **382**, eadd7046 (2023).
2. Phillips, J. W. *et al.* A repeated molecular architecture across thalamic pathways. *Nat. Neurosci.* **22**, 1925–1935 (2019).
3. Gao, C. *et al.* Molecular and spatial profiling of the paraventricular nucleus of the thalamus. *eLife* **12**, e81818 (2023).
4. Wallace, M. L. *et al.* Anatomical and single-cell transcriptional profiling of the murine habenular complex. *eLife* **9**, e51271 (2020).
